# LIPIDS MODULATE THE DYNAMICS OF GPCR:β-ARRESTIN INTERACTION

**DOI:** 10.1101/2024.03.16.585329

**Authors:** Antoniel Gomes, Michela Di Michele, Rita Ann Roessner, Marjorie Damian, Paulo M. Bisch, Nathalie Sibille, Maxime Louet, Jean-Louis Baneres, Nicolas Floquet

**Affiliations:** Institut des Biomolécules Max Mousseron(IBMM), UMR 5247 CNRS, Université de Montpellier, ENSCM, 1919 route de Mende, 34293 Montpellier cedex 5, France; Laboratório de Física Biológica, Instituto de Biofísica Carlos Chagas Filho, Universidade Federal do Rio de Janeiro, Brazil; Centre de Biologie Structurale (CBS), CNRS, Université de Montpellier, Inserm, Montpellier, France

**Author notes:** These authors contributed equally to this work (Antoniel Gomes, Michela Di Michele).

## Abstract

β-arrestins are key privileged molecular partners of G-Protein Coupled Receptors (GPCRs), triggering not only their desensitization but also intracellular signaling. Existing structures point to a high conformational plasticity of β-arrestin:GPCRs interaction, with two completely different orientations between receptor and β-arrestin. The same set of structures also indicates that the C-edge loop of β-arrestin could contribute to its anchoring to the membrane, through an interaction with specific lipids, namely PI(4,5)P2. Combining molecular dynamics simulations and fluorescence spectroscopy, we show that β-arrestin 1 interacts with membranes even in the absence of a receptor, an interaction that is enhanced by PI(4,5)P2 presumably holding the β-arrestin 1 C-edge loop into the lipid bilayer. This key interaction helps β-arrestin 1 to adopt a “receptor ready” orientation. As a consequence, PI(4,5)P2 also favors the coupling of β-arrestin 1 to the ghrelin receptor (GHSR). In addition, we show that β-arrestin can adopt the two known extreme orientations when complexed with GHSR. Of importance, PI(4,5)P2 shifts the equilibrium between the two different arrangements, favoring one of them. Simulations performed on the GHSR:β-arrestin complex suggest that release of the C-edge loop is required for these transitions to occur and point to a different distribution of PI(4,5)P2 around the complex depending on the orientation of receptor-bound arrestin. Taken together, our results highlight how PI(4,5)P2 plays a true third player role in the β-arrestin:GPCRs interaction, not only by preparing β-arrestin for its further interaction with receptors but also by modulating its orientation once the protein:protein complex is formed.

## Introduction

GPCRs form a large family of over 800 transmembrane receptors whose activities are linked to a wide range of physiological functions. As such, they constitute privileged targets for the design of new drugs ^1^. Their activities result from their coupling to intracellular effectors including G-proteins, GPCR kinases (GRKs) and arrestins that all initiate different signaling responses ^2^. In addition to GPCR desensitization, receptor-bound arrestins also initiate important signaling cascades, and thus regulate a wide range of cellular functions. Therefore, a better knowledge of arrestin/GPCRs interactions at the molecular scale is a key step for the design of novel drugs with biased signaling effects ^3^. Mammals express four different arrestins; while arrestin-1 (rod) and arrestin-4 (cone) are visual arrestins that bind exclusively to light-activated phosphorylated rhodopsin that are expressed mainly in the eyes, arrestin-2 (also called β-arrestin-1) and arrestin-3 (β-arrestin-2) are more ubiquitously expressed, regulating hundreds of different receptors ^4^. This already indicates a high degree of adaptability for β-arrestins which can thus recognize a large number of different receptors despite significantly different spatial distribution of amino acids and cellular environment. Available structures show that β-arrestins consist of two N- and C-terminal domains flanking a central region called the “finger loop” that inserts directly into the intracellular transmembrane bundle of GPCRs (**Figure 1A**). The N-domain of β-arrestin is dedicated to the binding of phosphorylated residues ^5,6^ usually located in the C-tail or intracellular loop-3 of receptors, again with a high degree of adaptability ^7^. The C-terminal domain plays a direct role in the interaction of β-arrestin with membranes; this interaction involves (1) a plausible insertion of its “C-edge loop” into the lipid bilayer ^8^, and (2) the binding of phosphatidylinositol-4,5-bisphosphates (PI(4,5)P2)^9^ and its derivatives ^10,11^ to a specific site. To date only 6 structures described β-arrestin complexed to different GPCRs, including rhodopsin (PDB:4ZWJ ^12^ & 5DGY ^13^ & 5W0P ^14^), beta-1 adrenergic (B1AR, PDB:6TKO ^15^), neurotensin-1 (NTS1R, PDB:6PWC ^16^ & 6UP7 ^9^), M2 muscarinic (PDB:6U1N ^17^), vasopressin 2 (PDB:7R0C ^18^) and serotonin 5-HT_2B_ (PDB:7SRS ^19^) receptors. Although limited, this set of structures already sheds light on the high degree of plasticity of β-arrestin:GPCR interactions. The overall orientation of β-arrestin in the protein:protein complex shows a quasi-rigid body rotation around 75° in the two NTS1R structures with respect to the M2, Beta-1, and visual rhodopsin structures (**Figure 1B**). In this respect, the two most recent Vasopressin 2 and 5-HT_2B_ receptor structures can be considered as intermediate orientations. Another key point concerns the putative insertion of the C-edge loop of β-arrestin in the membrane (**Figure 1C**), as suggested in some structures (mainly those describing PI(4,5)P2 bound to the C-domain). However, since most of these structures were obtained in detergents rather than in a lipid bilayer, it remains difficult to assess whether this loop is effectively inserted into the membrane under more physiological conditions. The last point of interest concerns the finger loop of β-arrestin, which is inserted into the transmembrane helix bundle of the receptor. Indeed, among the available structures the finger loop presents a wide diversity of conformations, different from those observed in isolated arrestins (**Figure 1D**). This is rather intriguing since this loop represents the main point of contact with the receptors, and as such one might expect a conserved binding mode throughout the whole GPCR family. Until now, these structural differences were mainly attributed to the nature of the coupled receptor and have been proposed to possibly help β-arrestin adapt to the different phosphorylation schemes of GPCRs. Here we combined molecular dynamics (MD) simulations and site-directed fluorescence spectroscopy with an isolated receptor in lipid nanodiscs to more thoroughly characterize the interaction of β-arrestin-1 with membranes in the absence or presence of the ghrelin receptor (GHSR), a prototypical class-A GPCR involved in growth hormone secretion and food intake ^20^ and metabolic homeostasis ^21^. By doing so, we reveal unprecedented features on the dynamics of the GPCR:arrestin arrangement and the impact of the membrane environment on these dynamics.

**Figure 1.**
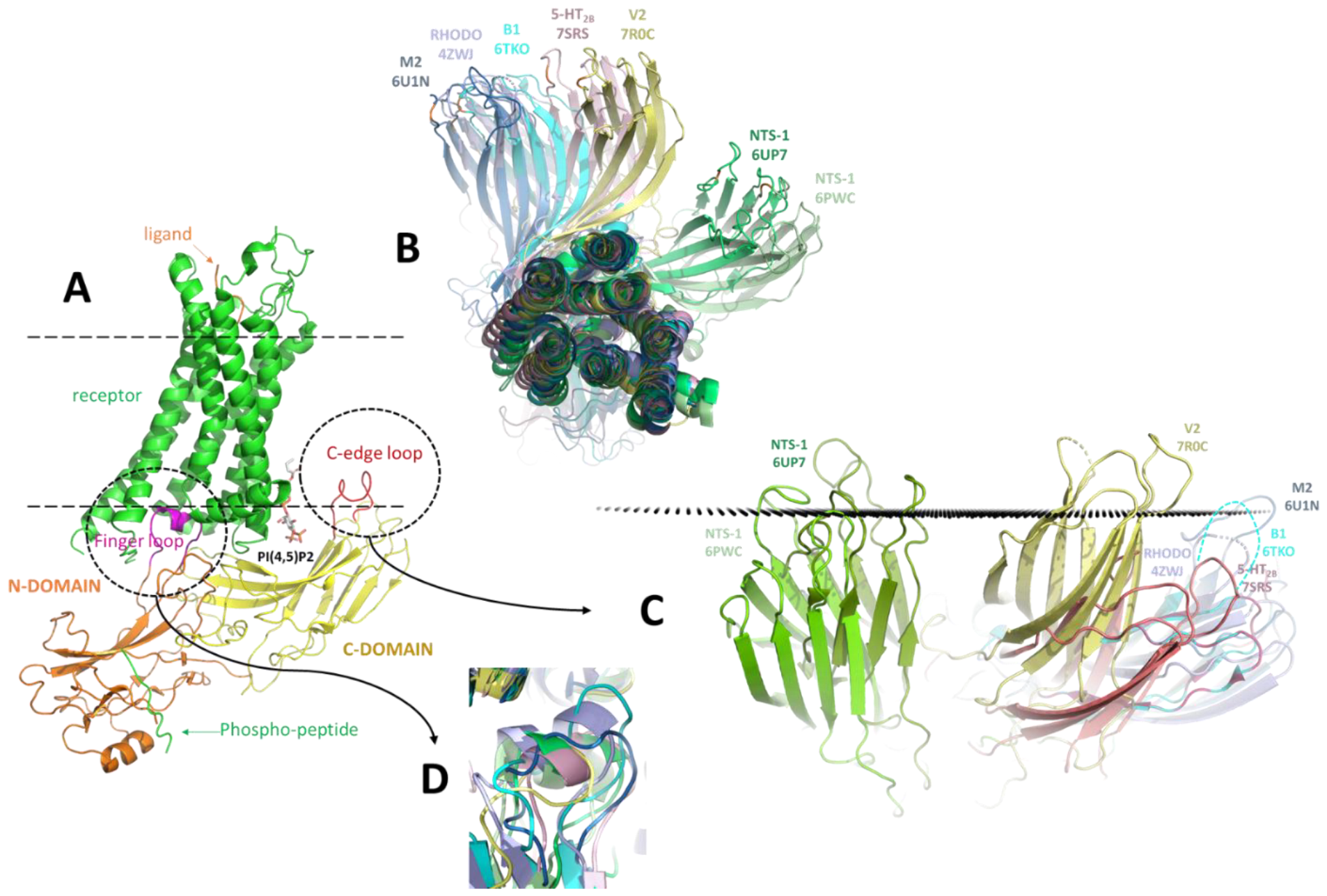
(A) Summary of β-arrestin contacts with the receptor and with the membrane as observed in the NTS1R structure (PDB:6UP7). (B) Orientations of β-arrestin in the different known structures of β-arrestin:GPCR complexes showing two main sets of orientations with NTS1R structures aside. (C) Position of the C-edge loop of β-arrestin in respect with the membrane among the same set of structures; black points correspond to the position of the membrane predicted by OPM (Orientations of Proteins in Membrane)^22^ for the complex with the beta-1 (B1AR) adrenergic receptor (PDB:6TKO). (D) Conformations adopted by the Finger-loop of β-arrestin in the different receptors.

## Results

### PI(4,5)P2 increases β-arrestin:membrane interactions

To analyze the ability of human β-arrestins to interact with membranes, a fluorescent probe, monobromobimane (MB), was inserted at positions 68 or 191 of a cysteine-free mutant of β-arrestin 1 (**Figure 2A**). The first modification (position 68, shown in orange in **Figure 2A**) has been extensively used in the field to probe GPCR:β-arrestin interactions, as changes in MB emission report on the insertion of the finger loop of β-arrestin in the receptor core ^23,24^. The second modification, with the probe at position 191 of β-arrestin 1 (reported in green) was designed to monitor the insertion of the C-edge loop in the lipid bilayer. This modification is similar to that used for analyzing the anchoring of visual arrestin to disk membranes in the absence or presence of rhodopsin ^25^. We then analyzed the emission properties of MB-labeled arrestin in the absence or presence of POPC/POPG nanodiscs with or without 2.5% PI(4,5)P2. Here we used 2.5% PI(4,5)P2 (lipid-to-PI(4,5)P2 molar ratio), as we previously established that this is the amount required for a maximal effect on GHSR functioning ^26^. In addition, spin-labeled PC (phosphatidylcholine) was included into the nanodiscs to trigger changes in MB emission upon insertion of the probe into the bilayer in the absence of receptor (see Experimental Procedures). In all cases, no change in the MB emission was observed when the probe was located in the finger loop of β-arrestin 1, indicating that this region remained distant from the membrane surface independently of the composition of the lipid nanodisc (**Figure 2B**). With regard to the probe at position 191, a significant change in the MB emission properties was observed for the nanodiscs containing PI(4,5)P2. This indicates that this region of β-arrestin 1 significantly interacts with the bilayer of the nanodisc, especially in the presence of this particular lipid.

**Figure 2:**
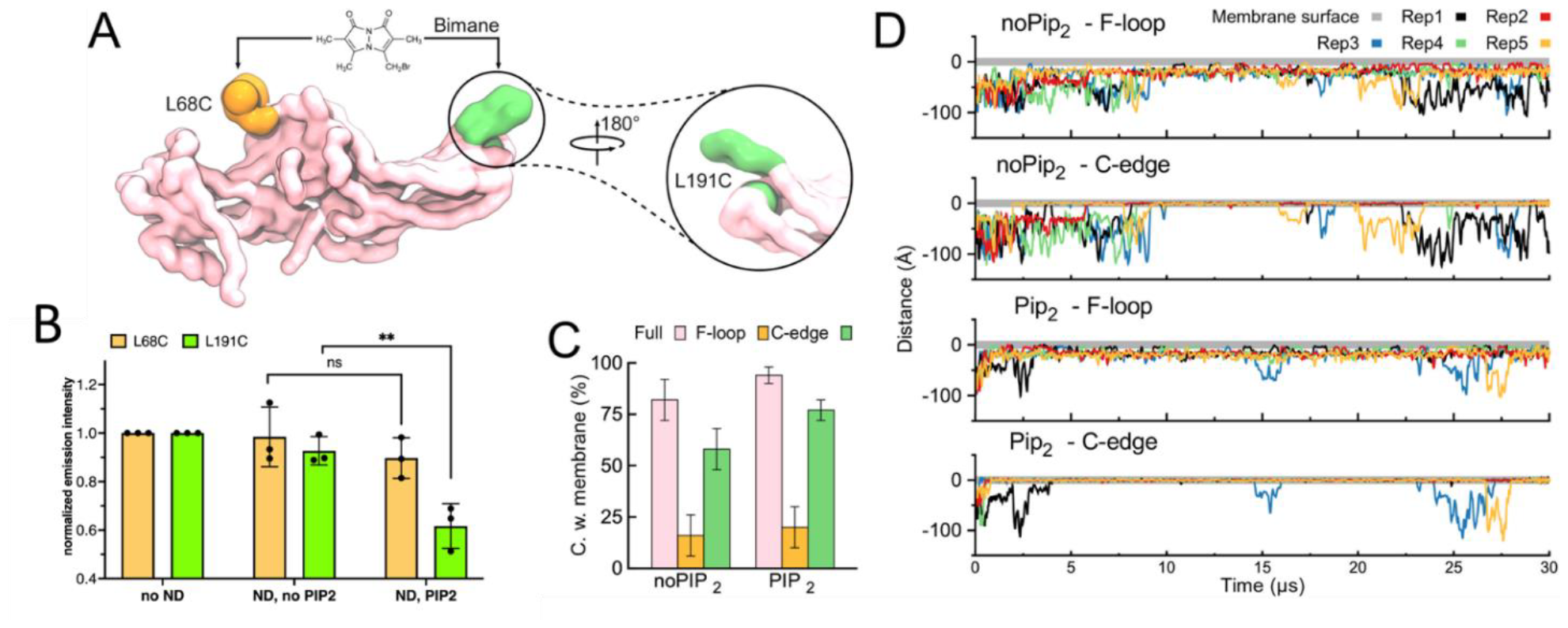
Structural aspects of β-arrestin 1 and its interaction with membranes. (A) Positions in β-arrestin 1 selected to insert MB. (B) Changes in the MB emission intensity for the two positions in the absence of nanodiscs (no ND), in the presence of POPC/POPG nanodiscs (ND, no PIP2) or in the presence of POPC/POPG nanodiscs containing 2.5% PI(4,5)P2 (ND, PIP2). Data are the mean value ± SD of three experiments. Statistical values were obtained by means of unpaired Student’s t-test (**p ≤ 0.01, ns: not significant). (C) Evaluation of contacts with the membrane made by the two L68 and L191 residues as well as by the full length β-arrestin during the CGMD simulations; bars correspond to the means and standard deviations calculated over five independent simulations. (D) Time-dependent analysis showing an increased insertion of the C-edge loop in the membrane in the presence of PI(4,5)P2 whereas the Finger-loop was always held apart from the surface. These plots also show that β-arrestin binds more quickly and more stably to the membrane containing PI(4,5)P2.

To get a molecular picture of this β-arrestin:membrane interaction, we then performed Coarse-Grained molecular dynamics (CGMD) simulations starting with β-arrestin 1 located at a minimal distance above 20 Å from the bilayer (*i*.*e*. a distance higher than the cutoff distance which was set to 12 Å) to prevent any initial bias. 5 simulations of 30 µs, were performed using the MARTINI 3 Force Field ^27^ and using a membrane composition very similar to that of the lipid nanodiscs, replacing or not 10 POPC lipids by PI(4,5)P2 in each leaflet aiming to capture the specific effect of this lipid. The simulations first confirmed the ability of the C-edge loop of β-arrestin to spontaneously insert into the lipid bilayer during 76.1%±6.3 and 55.4%±10.3 with/without PI(4,5)P2, respectively (**Figure 2C**). In contrast, position 68 was barely in contact with the membrane (19.7%±10.1 and 10%±5.7 with/without PI(4,5)P2 respectively) (**Figure 2C**), thus confirming the experimental results and indicating a probable tilt of β-arrestin’s orientation at the membrane surface. In contrast to the experiments, however, our simulations predicted a large proportion of β-arrestin at the membrane surface, even in the absence of PI(4,5)P2 (96.4%±2.7 and 83.4%±9.6 of the time with/without PI(4,5)P2, respectively) (**Figure 2C**). This over-estimation could plausibly be explained by the finite size of the simulation box that considerably increases the probability of interaction of the protein with the membrane, although unbinding events of β-arrestin to the membrane occurred during the simulations both with and without PI(4,5)P2 (**Figure 2D**). Our simulations further suggested a faster anchoring and a slower release of β-arrestin in the presence of PI(4,5)P2 (mean anchoring time ∼2.5 µs versus ∼9 µs without)(**Figure 2D**). Accordingly, we can reasonably anticipate that the interaction of the C-edge loop with the membrane would contribute to increase these differences on longer time-scales as those corresponding to the experiments.

Supporting the experiments, our computational results suggest a particular orientation of the β-arrestin at the membrane surface corresponding to an insertion of the C-edge loop into the bilayer and a position more distant from the surface for the finger-loop (**Figure 3A**). This orientation is consistent with the presence of PI(4,5)P2 in the previously identified cleft located in the C-domain of β-arrestin. To better understand how PI(4,5)P2 could contribute to the binding of β-arrestin to the membrane and to this specific orientation, we computed the spatial occupancy of the PI(4,5)P2 headgroups on the β-arrestin’s surface along all CG-MD trajectories. Interestingly, we retrieved an interaction hotspot for PI(4,5)P2 that exactly matches the region of the C-domain previously identified (**Figure 3B**). This region is enriched with positively charged residues, such as K250 and K324 previously identified as important binders for phosphatidyl-inositols ^9^. This emphasizes the ability of the MARTINI3 force field to predict successfully PI(4,5)P2 binding at the protein’s surface, as we already recently reported for the isolated ghrelin receptor ^26^. Though to a lesser extent PI(4,5)P2 was also observed at the bottom of the C-edge loop, indicating a second, less specific site for PI(4,5)P2, that possibly contributes to the retention of the C-edge loop in the bilayer after its anchoring. The same analysis performed on the other lipids not only confirmed that β-arrestin displayed a tilt angle at the membrane surface (**Figure 3C**), but also that PI(4,5)P2 could contribute to maintain the protein closer to the membrane surface, as previously observed in **Figure 2D**. Distributions of the “tilt” and “roll” angles of β-arrestin during our MD simulations (**Figure 3D**) clearly showed the same preferred orientation at the membrane surface of β-arrestin independently of the presence of PI(4,5)P2 and corresponding to restrained “tilt” values around 5-35° and “roll” values more largely distributed in the range 40-140°. Two subsets of orientations could indeed be defined from the obtained distributions corresponding to a “tilt” value of about 20° and to “roll” values of either 60° or 120°. The increased interaction of β-arrestin with the membrane in presence of PI(4,5)P2 contributed to enrich the populations of both sub-states with a slight preference for the one corresponding to the higher roll value (**Figure 3E**). “tilt” and “roll” angles calculated from existing structures of β-arrestin:GPCR complexes deposited in the Protein Data Bank interestingly yieldied similar values. Together, our results indicate that isolated β-arrestin presumably adopts an orientation at the membrane surface directly compatible with the interaction with a receptor, PI(4,5)P2 further contributing to increase the frequency of this orientation by stabilizing the position of the C-edge loop in the bilayer.

**Figure 3:**
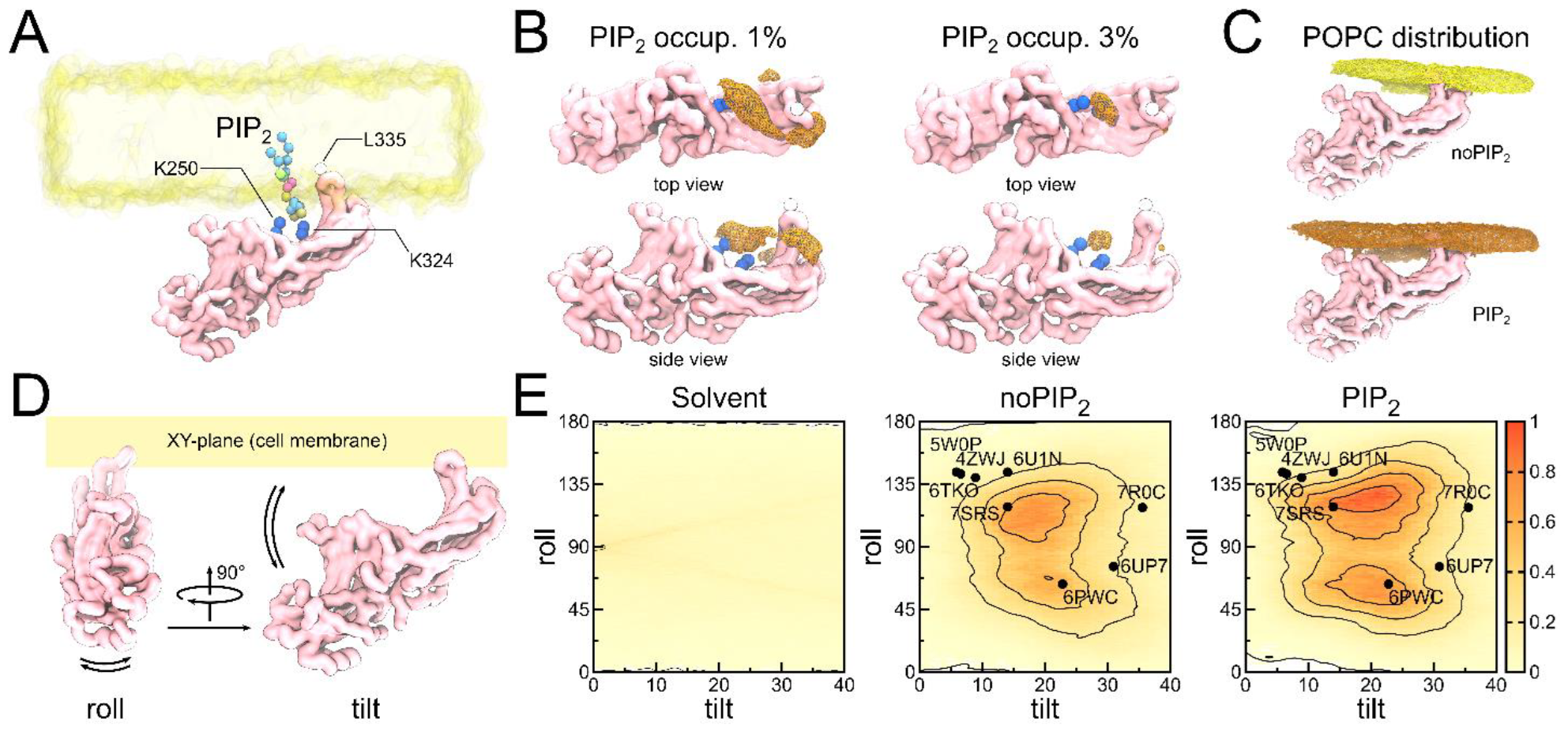
β-arrestin interaction with PI(4,5)P2 and predicted orientations to the cell membrane. (A) PI(4,5)P2 binding site as described in available structures. Important residues for membrane insertion of C-edge (L335), and PIP2 interaction (K250 and K324) are shown. (B) statistical distribution of PI(4,5)P2 (orange meshes) along the obtained CGMD trajectories. It confirmed the specific binding of PI(4,5)P2 in the same site as that described in the literature ^26^. A secondary site distributed around the C-edge loop was also found. (C) Mean position of β-arrestin at the membrane surface (represented here by the statistical distribution of POPC heads during the simulations performed in the absence or presence of PI(4,5)P2). (D) The definition of tilt and roll angles aimed to report the position of the protein as respect to the membrane surface. (E) β-arrestin assumes a specific orientation at the membrane surface (see lines contours), compatible with its further interaction with receptors. Black dots correspond to tilt/roll angles extrapolated from existing β-arrestin:GPCR structures in the PDB.

### PI(4,5)P2 favors the interaction of arrestin with GHSR

To further explore the role of PI(4,5)P2 in the β-arrestin:membrane:receptor interplay, the MB emission experiments were repeated in the presence of agonist-activated GHSR in the lipid nanodiscs. In a first stage, we used a C-terminally truncated pre-activated β-arrestin 1 (residues 1–382) ^28^, thus circumventing the requirement for receptor phosphorylation. As shown in **Figure 4A**, a significant change in the emission properties of MB attached to the β-arrestin 1 finger loop was observed with nanodiscs containing MK0677-loaded GHSR, whether PI(4,5)P2 was present or not in the nanodiscs, consistent with our previous reports ^20^. In contrast, the emission properties of MB attached to C191 changed only when PI(4,5)P2 was present, similarly to receptor-free nanodiscs. This points to a possible model where activated arrestin could interact with the receptor core independently of its interaction with the lipid bilayer, with a concomitant membrane insertion of the C-edge loop when PI(4,5)P2 is present. To assess whether this membrane insertion could favor the interaction of β-arrestin 1 with the ghrelin receptor, we repeated this experiment with wild-type β-arrestin 1. As shown in **Figure 4B**, a slight change in the emission properties of MB attached to C68 was observed, but only when the lipid nanodiscs contained PI(4,5)P2, *i*.*e*. under conditions where the C-edge loop inserts into the bilayer. Altogether, this suggests a possible model where insertion of the C-edge loop of arrestin into the lipid bilayer, favored by the presence of PI(4,5)P2, could favor the interaction of the finger loop with the cytoplasmic core of the agonist-activated receptor, overcoming, at least to some extent, the requirement of arrestin activation by the receptor phosphorylated C-terminal region.

**Figure 4:**
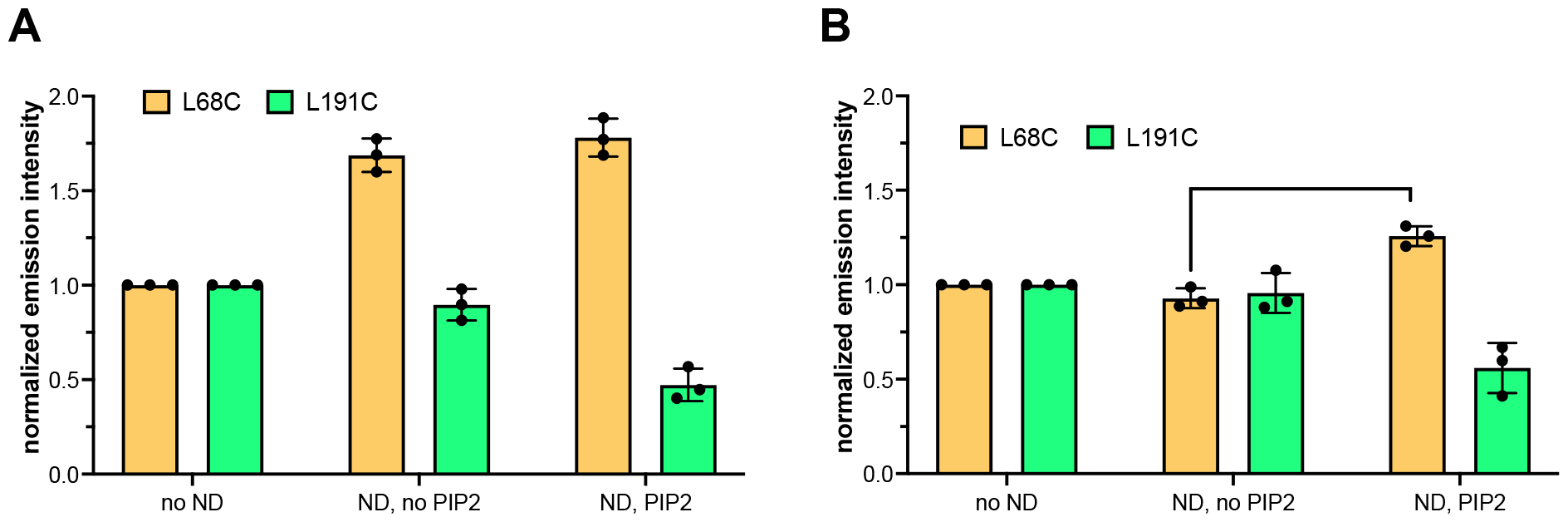
Role of the PI(4,5)P2 in the interaction of β-arrestin 1 with the ghrelin receptor. (A) Changes in the emission intensity of MB attached to either C68 or C191 of the pre-activated β-arrestin 1 mutant free in solution (no ND), in the presence of POPC/POPG nanodiscs containing unphosphorylated GHSR and 10 µM MK0677 (ND, no PIP2) or in the presence of POPC/POPG nanodiscs containing 2.5% PI(4,5)P2, unphosphorylated GHSR and 10 µM MK0677 (ND, PIP2). (B) Same experiment carried out with the wild-type β-arrestin 1 under the same conditions. In all cases, data are the mean value ± SD of three experiments. Statistical values in (B) were obtained by means of unpaired Student’s t-test (**p ≤ 0.01).

### PI(4,5)P2 promotes a specific orientation of β-arrestin in the complex with GHSR

Currently, it is not clear whether the insertion of β-arrestin 1 edge-loop in the bilayer caused by the presence of PI(4,5)P2 could contribute to select a specific orientation of this signaling protein with respect to the receptor it binds. Until recently, only the shifted structures of the NTS-1 receptor were indeed captured in the presence of PI(4,5)P2 ^9^ mimetics. However, the recently solved structures for the vasopressin 2 ^18^ and serotonin 5-HT_2B_ ^19^ receptors demonstrated that other (intermediate) orientations of β-arrestin are also compatible with the lipid bound to the same pocket. To address this key question, we built two homology models of the β-arrestin:GHSR complex using the two extreme orientations observed with the neurotensin NTS-1 (PDB:6UP7) and β1-adrenergic (PDB:6TKO) receptors as templates (**Figure 5**). Using these two models, we then selected several positions to insert fluorescent probes that would allow us to follow experimentally the orientation of β-arrestin with respect to GHSR. Namely, we selected F71^1.60^ for GHSR, as this residue had already been mutated to a single reactive cysteine and subsequently labeled with fluorescent probes in a previous analysis of the interaction of the isolated receptor in nanodisc and G proteins ^29^. For β-arrestin 1, we selected position 167 and 191, as they should report on the position of the receptor with respect to the arrestin N- and C-lobe. Moreover, these positions had already been modified for β-arrestin 1 intramolecular FRET experiments, and this has been shown not to affect the functional properties of this protein with regard to its interaction with GHSR in nanodiscs ^30^. Although the 71^1.60^:167 distance poorly discriminates the different orientations of β-arrestin in the receptor:arrestin complex (**Figure 5**), it nevertheless reports on the core engagement of β-arrestin with GHSR, and as such allows to rule out any possible tail-only engaged complex. On the contrary, the 71^1.60^:191 distance was much more informative to distinguish the different orientations of β-arrestin in the complex, as it is significantly different in structures of β-arrestin 1 complexes (about 34 Å in the complex with NTS1R and in the 55-63 Å range for the other receptors; see **Table 1**).

**Table 1:**
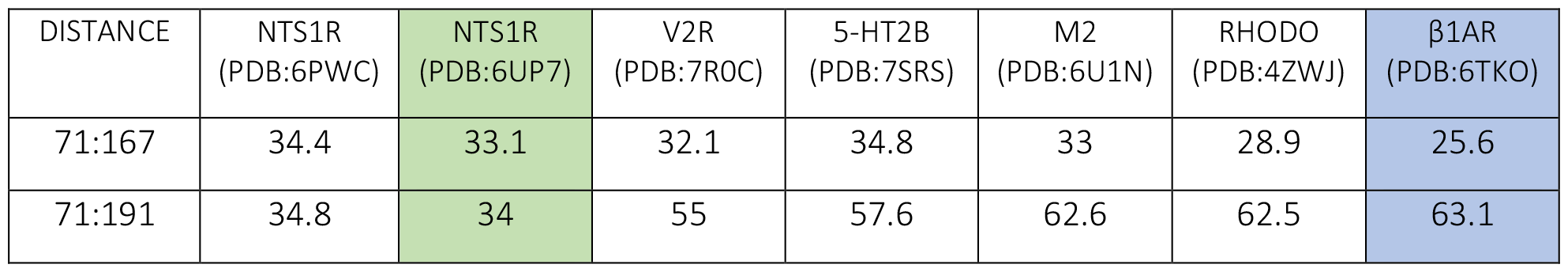
Intermolecular distances measured to discriminate between the different orientations of β-arrestin in its receptor-bound state. Distances between residue 71^1.60^ of GHSR and either residues 167 or 191 of β-arrestin 1, as measured in all available β-arrestin:GPCR complexes in the PDB.

**Figure 5:**
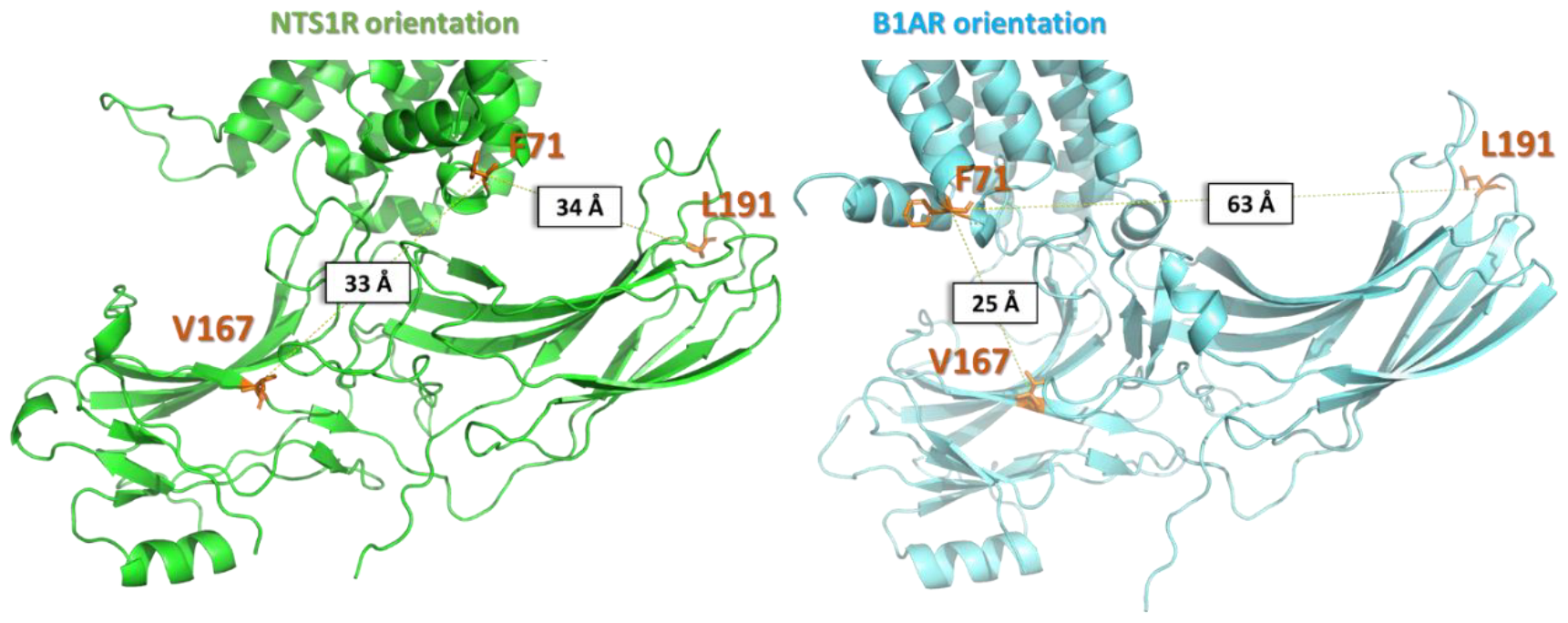
Models of the β-arrestin:GHSR complex built from NTS1R (green, PDB:6UP7) or β1AR (blue, PDB:6TKO) :β-arrestin 1 structures. The location of the residues used to attach the fluorescent probes in β-arrestin 1 (V167 or L191) and in GHSR (F71^1.60^) are depicted in orange sticks.

We thus analyzed the interaction between MK0677-activated GHSR and β-arrestin 1 by monitoring the LRET signal between the receptor labeled with Alexa Fluor 488 or BodipyTMR at position 71^1.60^ and β-arrestin 1 labeled with a Tb cryptate donor either at position 167 or at position 191. For the 71^1.60^:167 pair, Alexa Fluor 488 was used as the fluorescence acceptor, as the R_0_ value with the Tb cryptate as the donor, i.e. 39 Å, is in the range of the expected distance values (**Table 1**). For the 71^1.60^:191 pair, BodipyTMR was used as the acceptor as it displays a higher R_0_ value (50.8 Å) compatible with distances in the 40-60 Å range. We used LRET, as this technique has several technical advantages over conventional FRET, including distance measurement with greater accuracy, and insensitivity to incomplete labeling ^31^. To circumvent the possible bias due to the use of an unphosphorylated receptor and a pre-activated arrestin, we instead used sortase to enzymatically ligate to the GHSR transmembrane core a synthetic peptide where the potential phosphorylation sites in the GHSR C-terminal domain ^32^ had been replaced by their phosphorylated counterpart (see Experimental Procedures). Of importance, the interaction pattern observed with this receptor variant and wild-type β-arrestin 1 was similar to that obtained with the wild-type ghrelin receptor phosphorylated *in vitro* with recombinant GRK5 (**supplementary Figure S1**), fully validating our strategy.

Although low amounts of purified protein were obtained using this ligation approach, this strategy ensured homogeneous phosphorylation of the receptor compared to *in vitro* phosphorylation with recombinant GRK, justifying its use in the present experiments. The LRET profiles were thus recorded with this modified GHSR inserted into nanodiscs with or without 2.5% PI(4,5)P2. In all cases, these profiles were best fitted with a multi-exponential, with two major components, a slow and a fast one (**Figure 6**).

**Figure 6:**
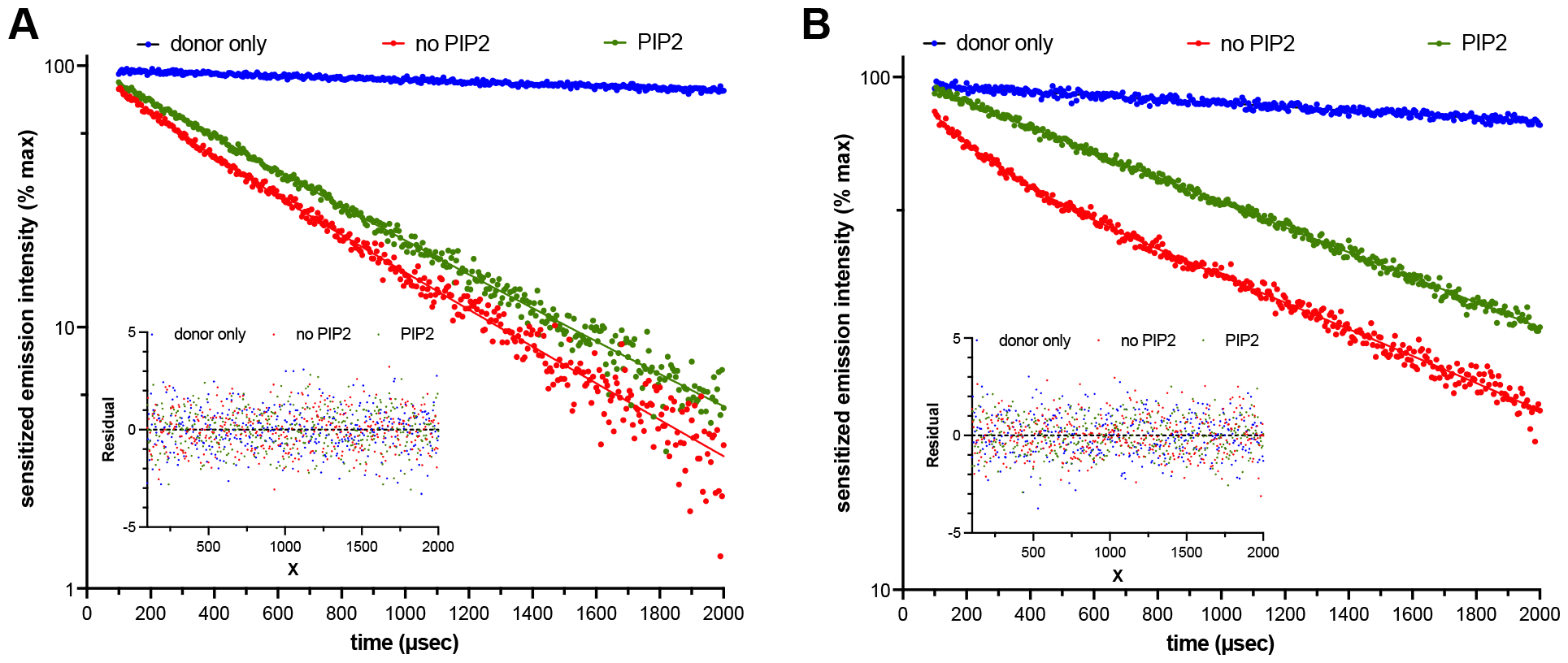
Relative orientation of the protein components in the GHSR:arrestin complex. Sensitized-emission decays from GHSR labeled with Alexa Fluor 488 (A) or Bodipy-TMR (B) on C71^1.60^ and assembled in nanodiscs containing or not 2.5% PI(4,5)P2, in the presence of β-arrestin 1 labeled either on C167 (A) or on C191 (B) with Lumi-4 Tb, and of 10 μM MK0677. Data are presented as normalized fluorescence intensity at 515 nm as a function of time and represents the average of three measurements. Residual values represent the goodness of the multi-exponential fits.

The distance between the LRET donor and acceptor was then approximated from the sensitized-emission lifetimes of these two components, and their relative populations were calculated from the pre-exponential factors and the excited state lifetime values ^33^. As shown in **Table 2**, the emission decays corresponded to a similar distance between the probes independently of the presence of PI(4,5)P2 in the nanodisc. These were in the same range as those estimated for the major orientations observed in the cryoEM structures (report to **Figure 5 and Table 1**), indicating that both orientations could co-exist in solution. However, the relative population of each arrangement was highly dependent on the presence of PI(4,5)P2 in the nanodiscs. Indeed, whereas both populations were almost similar in the absence of PI(4,5)P2, the NTS1R-like arrangement was significantly under-represented when PI(4,5)P2 was present in the bilayer, suggesting that interaction of β-arrestin 1 with the lipid bilayer in the nanodisc, as evidenced in the MB-emission assays, could favor one of the possible geometrical arrangements of the complex, namely the β1AR-like one.

**Table 2:** Approximative distance (D) between the fluorescent probes calculated using the sensitized-emission lifetimes obtained from the two dominant exponential components of the sensitized-emission decays in Figure 6. The molecular fractions A1 and A2, expressed in % of the total population, were calculated from the pre-exponential factors and the excited state lifetime values.

|  | No PIP2 |  |  |  | PIP2 |  |  |  |
| --- | --- | --- | --- | --- | --- | --- | --- | --- |
|  | D1 (Å) | A1 (%) | D2 (Å) | A2 (%) | D1 (Å) | A1 (%) | D2 (Å) | A2 (%) |
| 71-167 | 27.2±0.8 | 48.1±5.6 | 33.4±1.2 | 51.9±4.4 | 28.8±0.9 | 17.4±4.9 | 34.1±1.3 | 82.6±5.2 |
| 71-191 | 33.5±1.1 | 43.1±4.3 | 58.5±0.7 | 56.9±3.8 | 32.1±1.2 | 13.2±4.7 | 60.7±1.1 | 86.8±5.2 |

To get a molecular view of how PI(4,5)P2 could contribute to select this specific orientation of β-arrestin in the complex with GHSR, we performed coarse-grained MD simulations starting from the two extreme orientations already depicted in **Figure 5**. A standard elastic network was defined as classically done in studies in which the MARTINI force field is used with some minor modifications: first we suppressed the springs involving residues of the finger loop so that the latter could adapt to the receptor, as shown in available structures (**Figure 1D**). The same was done for residues of the intra-cellular loop 3 of the receptor, which is not seen in existing GHSR structures, indicative of its high flexibility. It must be noted here that in both NTS1R-like and β1AR-like starting models, the C-edge loop of β-arrestin was sufficiently high to be initially inserted in the lipid bilayer. Finally, a restraint was added to the finger loop so that it was maintained inside the receptor core during the simulations, thus limiting any release of β-arrestin from the membrane. 20 simulations of 30 µs were obtained starting from both orientations and in the presence or absence of PI(4,5)P2 (four different systems corresponding to a total simulation amount of 2.4 ms). To follow the orientation of β-arrestin with respect to the receptor, we defined a rotation angle using the *x,y* components of the vector defined by residues 71 of the receptor and 191 of β-arrestin (after alignment onto the receptor’s backbone beads) and taking the NTS1R orientation as a reference. This way, values around 0 indicate an NTS1R orientation whereas values around 75° correspond to B1AR orientation. The distributions reported in **Figure 7A** indicated that no spontaneous transition was observed from the B1AR to the NTS1R arrangement, whether PI(4,5)P2 was present in the membrane or not. On the contrary, starting from the NTS1R orientation, several transitions towards the B1AR orientation were observed, regardless of the presence of PI(4,5)P2. These results showed that the transition between the two orientations can be observed without the release of β-arrestin from the membrane. The 2D plot of **Figure 7B** computed from simulations starting from the NTS1R orientation interestingly showed a good correlation between the β-arrestin rotation angle and the 71:191 distance, with two main populations centered around 34 and 60 Å respectively, in agreement with the values measured in the experiments.

**Figure 7:**
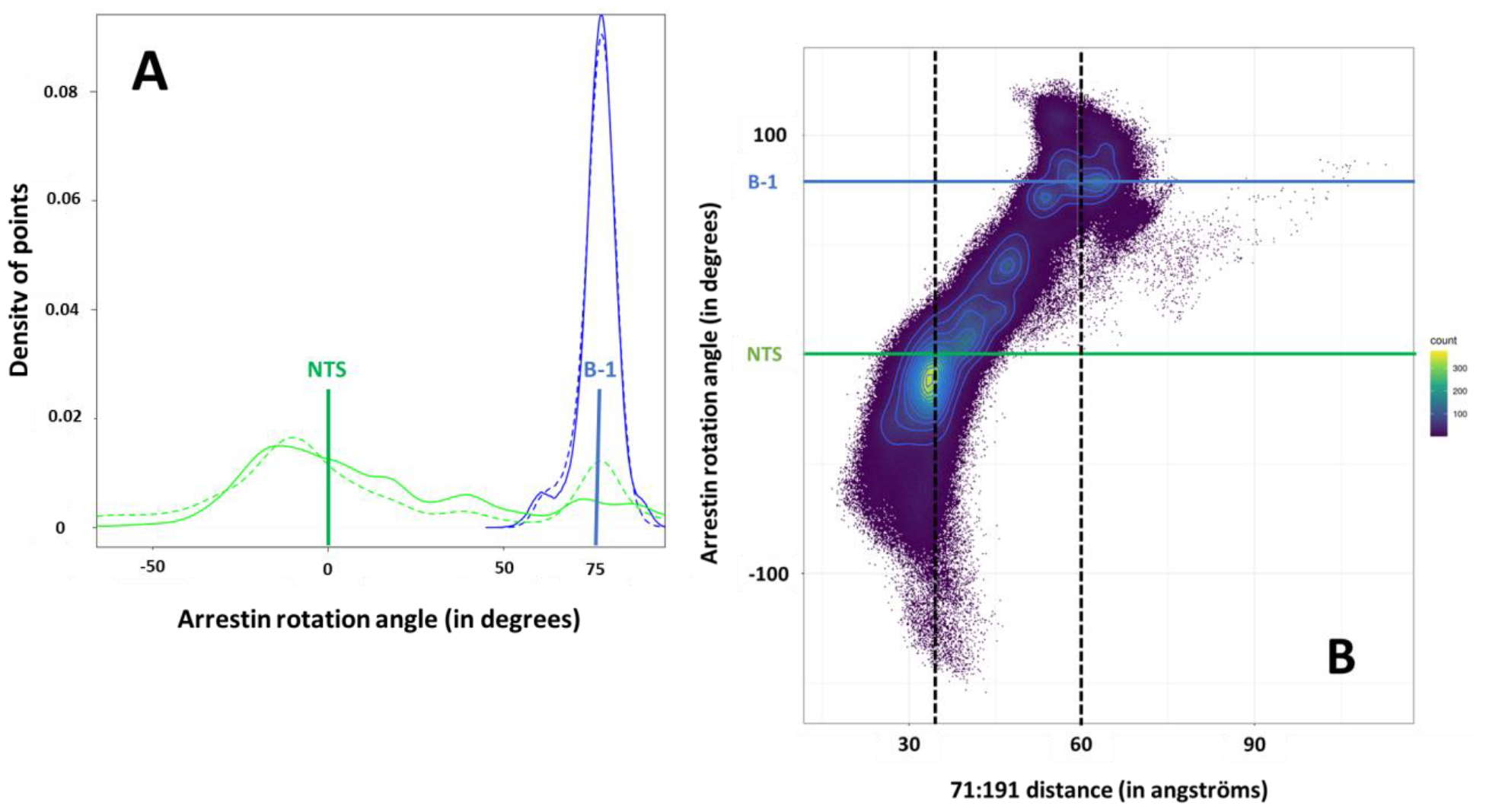
Results from CG simulations of the GHSR: β-arrestin complex starting from the NTS1R (green) or B1AR (blue) orientations. (A) Distribution of the rotation angle of β-arrestin among all CG simulations performed with (full line) or without (dashed line) PI(4,5)P2. (B) 2D plot corresponding to data obtained starting from the NTS1R orientation and showing the correlation between the β-arrestin rotation angle and the 71:191 distance that was measured experimentally.

### C-edge loop release appears as crucial for transitions between both orientations

Consequently, we further used the 71:191 distance as a collective variable for meta-dynamics simulations aiming to promote transitions in both directions (not only from NTS1R to B1AR as already observed in free CG MD but also from B1AR to NTS1R). All the simulations were stopped after ∼15 µs. Overall, the free energy profiles resulting from these CG meta-dynamics simulations confirmed the existence of two equivalent minima compatible with the measured distances and corresponding to the two extreme orientations of β-arrestin (**supplementary Figure S2 A**,**B**,**C&D**). As already observed in the free MD simulations, transitions from NTS1R to B1AR appeared easier than those from B1AR to NTS1R, although transitions from B1AR to NTS1R were nevertheless captured. The presence of (PI4,5)P2 seems to decrease the number of transition events. Accordingly, and as done above for the free MD simulations, only the meta-dynamics simulations sampling a high number of transitions (*i*.*e*. starting from NTS1R orientation) were used for further analysis. We observed multiple back and forth transitions between the two orientations in these simulations (**supplementary Figure S3**). All the conformations obtained were separated into four different subsets : (1) those corresponding to the NTS1R (0±10°) or (2) to the B1AR (70±10°) orientation, as well as those corresponding to transitions from (3) B1AR towards NTS1R or (4) NTS1R towards B1AR (**supplementary Figure S3**).

Looking at the main position of the C-edge loop in these subsets of conformations, we can first conclude that its interaction with the membrane increases in the presence of PI(4,5)P2, independently of the orientation of β-arrestin (**Figure 8A**). The C-edge loop was more often attached to the membrane in the B1AR orientation whereas it was more detached in the NTS1R orientation as well as in conformations corresponding to transitions (from B1AR to NTS1R or from NTS1R to B1AR). This was somewhat unexpected as β-arrestin was more tilted in the NTS1R starting orientation with a C-edge loop inserted deeper into the membrane. In all our simulations however, a downward movement of the loop was observed to reach a position relative to the membrane close to that observed in the B1AR orientation. Computing the probabilities of PI(4,5)P2 contacts around the complex showed significant differences which could explain a preference for the B1AR orientation in the presence of PI(4,5)P2 (**Figure 8B**). First, an additional PI(4,5)P2 binding site was observed at the interface between TM6 and TM7; this site was of particular interest as we already suggested that the binding of PI(4,5)P2 in this region could contribute to maintain an active conformation of the GHSR ^26^. Second, an increased presence of PI(4,5)P2 was noted at the bottom of the C-edge loop in the same site we already observed for the isolated β-arrestin (report to **Figure 3B** for comparison). Taken together, our results suggest that reducing the C-edge loop release from the membrane PI(4,5)P2 could promote the B1AR orientation (1) because of a preference for an anchored C-edge loop in this orientation, (2) because of the requirement for C-edge loop release to transition from B1AR to NTS1R and (3) because of additional binding sites of PI(4,5)P2 around the complex in the B1AR orientation, including that at the base of the loop that helps to maintain the loop into the bilayer. To confirm this possibility of a different distribution of PI(4,5)P2 around the complex according to the orientation of β-arrestin, we performed all-atoms simulations starting from the same models with three different replicas for each system (B1AR/NTS1R, PIP2/noPIP2), each lasting 2 µs. No transition between the two orientations was observed in the related trajectories (**supplementary Figure S4A**). Nevertheless, we observed a distribution of PI(4,5)P2 around the complex that was closely related to that captured in the CG simulations, including the sites 1, 2 & 3 we already described for the isolated receptor and the additional site at the bottom of the C-edge loop of β-arrestin (**supplementary Figure S4 B+C+D**). As observed in the CG simulations, an increased presence of PI(4,5)P2 was captured in site 3 when β-arrestin was in the B1AR orientation.

**Figure 8:**
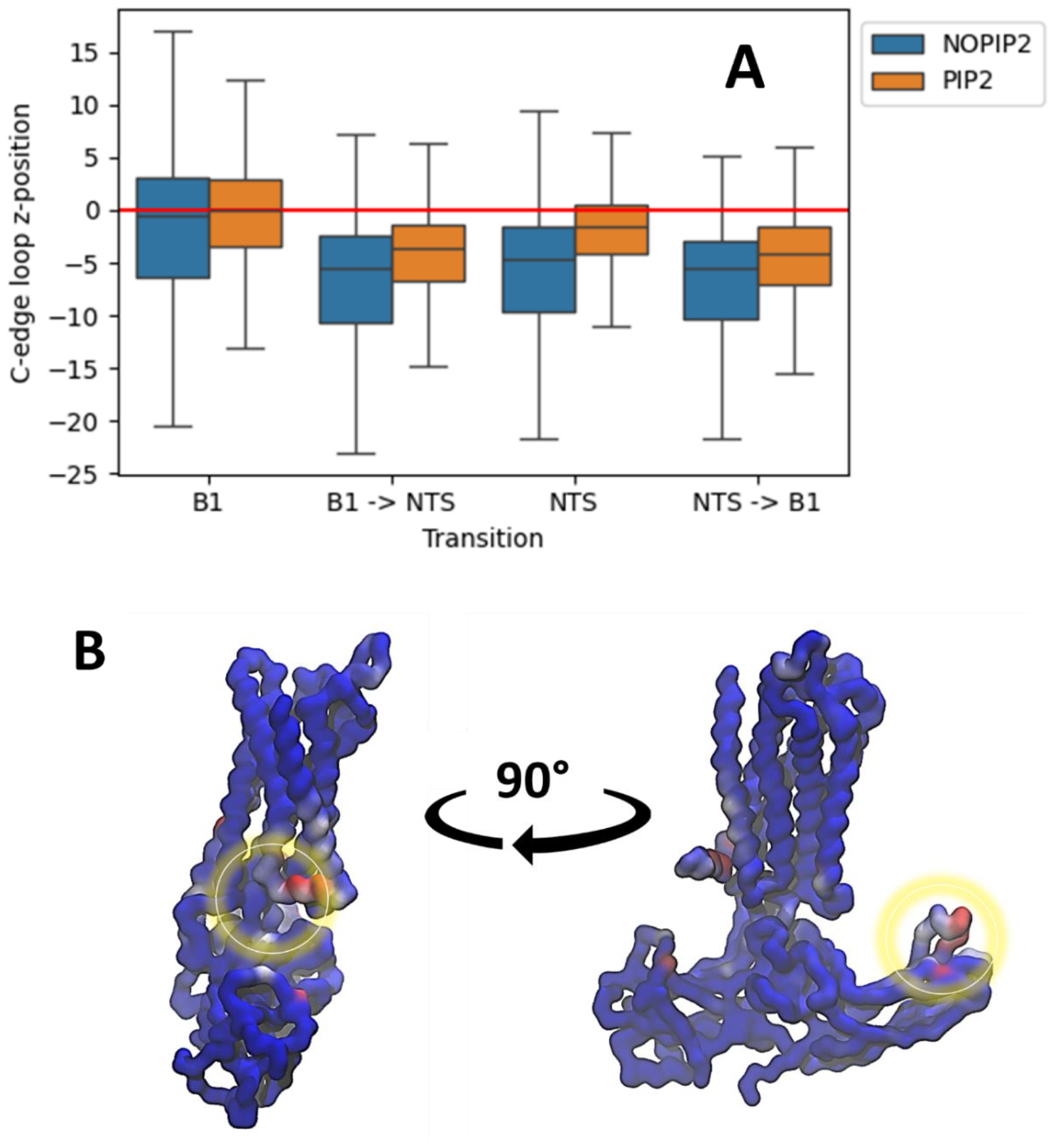
(A) Statistical analyses of the *z*-position of the C-edge loop respectively to the membrane in the four subsets of snapshots corresponding to the B1AR or NTS1R orientation or transitions between the two states; only the trajectories starting from NTS1R orientation were considered in this analysis. The red line corresponds to the position of the membrane. (B) contacts of PI(4,5)P2 around the GHSR: β-arrestin complex, focusing on regions that are more occupied in the B1AR orientation than in the NTS1R one. The color scale corresponds to an increase of 0 (blue) to 20% (red).

## DISCUSSION

There is increasing evidence that the membrane environment influences the conformational dynamics and pharmacological properties of GPCRs. In this context, we have used computational and fluorescence approaches to provide evidence for a direct interaction of β-arrestin 1 with the membrane, which is modulated by a specific lipid moiety, PI(4,5)P2. In addition, we showed that the different orientations of β-arrestin 1 captured in the different cryoEM structures of stabilized complexes with GPCRs can coexist in solution for the same receptor. This suggests that GPCR:arrestin coupling is a dynamic process that may involve multiple assemblies. Additionally, the membrane environment influenced the arrangement of the complex, as the presence of PI(4,5)P2 significantly affected the relative distribution of the different states. Taken together, our results pave the way for a detailed understanding of the molecular processes underlying the allosteric modulation of GPCRs by PI(4,5)P2 and, by extension, other lipids. We recently demonstrated that PI(4,5)P2 is directly involved in the activation of GHSR and its ability to further activate its G protein partner. Here, we provide evidence that PI(4,5)P2 not only affects GHSR-catalyzed G protein activation but also arrestin recruitment. Indeed, our data indicate that PI(4,5)P2 favors β-arrestin 1 coupling to the ghrelin receptor inserted into lipid nanodiscs, in agreement with previous data reporting a similar effect, especially for GPCRs that transiently interact with β-arrestins ^34^, which is the case of GHSR. Of importance, besides the recruitment to the agonist-activated receptor, we also observed a direct interaction of β-arrestin 1 with the membrane independent of GHSR. Again, this interaction was further modulated by PI(4,5)P2. This suggests that β-arrestin could operate at the membrane independently of its activating receptor, in line with previous observations ^35^. However, whether in our system this interaction with the lipid bilayer affects the conformational equilibria of β-arrestin is still an open question. In all cases, *i*.*e*. in the absence or presence of the receptor, our data point to the C-edge loop as a critical element in the β-arrestin:membrane interplay, consistent with previous MD simulation reports ^24,36^. Hence, this C-edge loop appears as a central element in β-arrestin coupling and activation, certainly cooperating with other structural elements (finger loop, phosphate sensor, polar core…) to achieve the final signaling pattern. Importantly, the orientation adopted by β-arrestin 1 at the bilayer surface in the absence of GHSR was closely related to that it adopts in the different complexes with GPCRs reported to date. This suggests a model in which the interaction with the bilayer would prepare the β-arrestin 1 protein for its recruitment to the activated receptor. If this is the case, then pre-association of β-arrestins with the membrane would play a critical role in β-arrestin 1 function by facilitating its interaction with the receptor and subsequent activation. This would be consistent with recent single-molecule microscopy observations in which β-arrestin spontaneously pre-associates with the lipid bilayer, allowing it to diffuse laterally until it encounters activated receptors ^36^. A few cryoEM structures of GPCR:β-arrestin 1 complexes have been solved so far. Notably, these structures report different orientations of β-arrestin relative to the receptor. One possibility would be that these different orientations depend on the conditions used to obtain these structures (sample engineering, receptor phosphorylation, complex preparation, detergents versus lipid bilayers…). Alternatively, it has been proposed that these orientations are receptor specific, in particular related to the presence of a phosphorylated ICL3 or a short C-tail on the GPCR ^16^. The data we obtained here with GHSR indicate that both arrangements can coexist for the same receptor. This suggests that GPCR:β-arrestin coupling is a dynamic process, consistent with recent cross-linking experiments ^37^. Importantly, the distribution of different states was here directly influenced by the nature of the lipid environment surrounding the receptor, namely the presence of PI(4,5)P2 molecules bound to specific sites. One possibility would therefore be that, for the same GPCR, different arrangements of the receptor:β-arrestin complex could occur. The relative population of these states would depend on the environment, here the lipids, and possibly also on other features such as the phosphorylation status of the receptor (the so-called phospho-barcode). Although we have no experimental and/or computational evidences at this stage of the analysis, it is tempting to then speculate that these different arrangements of the complex could be associated with (subtly) different conformational states of β-arrestin, with direct consequences for its functional properties. Whatever the details, our data thus indicate that GPCR:β-arrestin coupling is an inherently dynamic process under the control of multiple inputs, among them the membrane environment. As such, the latter appears as an integral player of this cooperative, dynamic interplay that ultimately leads to the β-arrestin activation and signaling patterns.

## Methods

### Modelling

#### Initial structures

The initial conformation of β-arrestin was modeled from its inactive structure available in the PDB (PDB:1G4M)^38^. The Modeller v9.19 ^39^ was used to fill missing residues of that protein, selecting the model displaying the lowest DOPE score value for the following simulations.

As no structure is yet solved for the β-arrestin:GHSR complex, initial models were generated by fitting the GHSR active conformation (PDB:7F9Y ^40^) onto the BETA-1 or the NTS-1 receptors complexed to β-arrestin (PDBs:6TKO ^15^, 6PWC ^16^). Doing so, no initial steric clash was observed. The Ghrelin peptide was removed from the resulting complexes before running the simulations.

#### AA simulations

All-Atoms (AA) MD simulations were performed with Gromacs 2020.4 ^41^ on either isolated β-arrestin in water or on the β-arrestin:GHSR complex inserted in a lipid bilayer. AA MD simulations were all carried out using the CHARMM36M force-field ^42^. The different systems were set up with tools available in CHARMM-GUI ^43^. Their energy was first minimized through 500,000 steps of the steepest descent algorithm using a convergence criterion of 1,000 kJ.mol^-1^.nm^-1^ and a switching function handling Van der Walls forces in the 10:12 Å range. Energy minimization was followed by successive equilibration phases in the NVT ensemble each lasting 25 ps, using an integration step of 1 fs and a Nosé-Hoover thermostat to maintain the temperature at 300K. Several replicas were run using different seeds therefore assuming a random assignment of initial velocities. Long-range electrostatic interactions were computed using the Particle Mesh Ewald (PME) algorithm. During the minimization and the equilibration processes heavy-atoms of the protein were restrained in position with force constants of 400.0 kJ.mol^-1^.nm^-1^ and 40.0 kJ.mol^-1^.nm^-1^ respectively applied to backbone and side-chains heavy atoms. The subsequent production runs were performed in the NPT ensemble with an integration step of 2 fs maintaining pressure to 1 bar through a Parrinello-Rahman barostat. For the different systems, the resulting MD trajectories from the production runs were all concatenated and analyzed as a whole.

#### CG simulations

Simulations at the Coarse-Grained (CG) level were performed using Gromacs ^44^ version 2020.4 and the MARTINI3 force field ^27^. β-arrestin and β-arrestin:GHSR complexes were first converted to MARTINI3 including an elastic network using the martinize2 script. The membrane part was generated using the insane script provided with MARTINI tools. The membrane composition was set to 48% of POPC, 32% of POPG, and 20% of cholesterol in each lipid layer to fit to that used in the experiments. To study the effect of PI(4,5)P2, a second membrane was built in which 10 POPC were replaced by 10 PI(4,5)P2 molecules in each leaflet. Parameters for PI(4,5)P2 were kindly provided by the group of Paulo Telles de Souza. Isolated β-arrestin was placed at an initial distance above 20 Å from the membrane surface to prevent any initial bias whereas the initial orientation to the membrane of the β-arrestin:GHSR complex was chosen according to the OPM method ^22^. After solvation, the global charges of the different systems were neutralized by adding Na^+^ and Cl^-^ ions to reach a salt concentration of 0.15 M. Energy minimization and equilibration steps were done according to the CHARMM-GUI protocol. During equilibration, the positional restraints applied to proteins (backbone atoms) or to lipids (heads) were progressively released whereas the integration step was increased from 2 to 20 fs. Initial velocities were randomly generated following a Maxwell-Boltzmann distribution at 300 K, maintaining the temperature using the V-Rescale thermostat. A semi-isotropic pressure coupling was applied maintaining the pressure at 1 bar using the Berendsen barostat. Production runs were performed using the Parrinello-Rahman barostat and all stopped after reaching ∼30 µs.

#### Meta-dynamics

Well-tempered metadynamics simulations were performed at the CG level using PLUMED ^45^ (version 2.8) employing the same parameters as those used in free MD and a biasfactor value of 10. The chosen collective variable was the inter-residue distance that was measured experimentally (*i*.*e*. between F71 and L191 residues) with no wall applied so that the whole phase space was accessible during the simulations. Gaussians of 0.2 nm width and 0.2 kJ.mol^-1^ height were deposited every 5000 steps (100 ps). These parameters allowed us to sample the expected range of distances in µs time-scale simulations. Simulations were stopped after ∼15µs.

#### Optimizing EN for isolated β-arrestin

As the elastic network (EN) has a strong influence on protein dynamics ^46^, we generated and tested the ability of different EN models to reproduce AA simulations data obtained on isolated β-arrestin. These data included the populations of the most representative conformations (clusters), the RMSF values computed on the C_α_ and C_α_-C_α_ covariance matrix, see **Table 3**.

**Table 3:** data resulting from CG simulations performed on isolated β-arrestin and validating the choice of a modified EN model.

| Positional covariance cut-off | Default Enledyn | 0.0 | 0.1 | 0.2 | 0.3 | 0.4 | 0.5 | 0.6 | 0.7 | 0.8 |
| --- | --- | --- | --- | --- | --- | --- | --- | --- | --- | --- |
| Common conformers (%) | 77.93 | 89.29 | 86.70 | 92.67 | 81.65 | 68.28 | 82.85 | 11.58 | 10.43 | 10.34 |
| RMSF correlation coefficient | 0.816 | 0.943 | 0.926 | 0.940 | 0.948 | 0.911 | 0.935 | 0.914 | 0.843 | 0.774 |
| Covariance matrix correlation coefficient | 0.909 | 0.928 | 0.931 | 0.940 | 0.935 | 0.942 | 0.930 | 0.836 | 0.901 | 0.779 |

EN models were generated from 6 independent replicas of 500 ns performed on isolated β-arrestin at the AA level. For this, we downscaled the spring force constant from the standard 500 kJ.mol^-1^.nm^-1^ of elnedyn to 0 kJ.mol^-1^.nm^-1^ in respect to residue-pair distance variance observed during all-atom simulations (see equation below), inspired by other work ^47^.

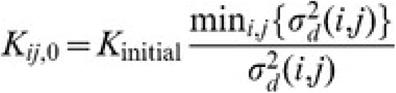

where K_ij_ is the new force constant for the residue-pair of residue i and j, K_initial_ being the initial force constant set up at 500 kJ.mol^-1^.nm^-1^ (as in the standard Elnedyn elastic network), σ_d_(i,j) is the variance of the distance for the residue-pair i j. Finally, min{σ_d_(i,j)} is the minimal variance over all possible residue pairs. With this equation, the elastic of the residue-pair distance which varied less is set up to K_initial_, while all other force constants are scaled down inversely proportional to the fluctuations of the inter-residue distance.

In addition we deleted springs when the positional covariance was below a certain cutoff and/or the mean distance along the all-atom trajectory was above 9.0 Å. We tested different cutoffs for the positional covariance from 0.0 to 0.8 (incremented by 0.1). A cutoff for the positional covariance of 0.2 was found to be the best network model which best reproduced the all-atom behavior with respect to the three metrics described above. This model was retained for the CG simulations of isolated β-arrestin, whereas a classical EN was used for simulations performed on the GHSR: β-arrestin complex.

### Experiments

#### Protein preparation

Human GHSR and its mutants were expressed in *E. coli* inclusion bodies, folded in amphipol A8-35 from an SDS-unfolded state and then A8-35 was exchanged to n-Dodecyl-β-D-Maltopyranoside (β-DDM) as described ^20^. For reconstitution into nanodiscs, the His-tagged receptor in a 50 mM Tris-HCl pH 8, 100 mM NaCl, 2 mM β-DDM was first batch-bound onto a pre-equilibrated Ni-NTA superflow resin. The slurry was then mixed with 10 µM of JMV3011, MSP1E3D1(-) and a POPC:POPG (3:2 molar ratio) mixture containing or not 2.5% PI(4,5)P2 molar ratio at a 0.1:1:75 receptor:MSP:lipid ratio, with the receptor still immobilized on the Ni-NTA matrix. After 1h incubation at 4°C, polystyrene beads (Bio-Beads SM-2) were added at an 80% (w/v) ratio and incubated under smooth stirring for 4 hours at 4°C. The resin was then extensively washed with a 50 mM Tris-HCl pH 8, 150 mM NaCl, 1 µM JMV3011 buffer and the His-tagged receptor eluted with the same buffer containing 200 mM imidazole. After extensive dialysis in a 25 mM HEPES, 150 mM NaCl, 0.5 mM EDTA, pH 7.4 buffer, active receptor fractions were purified using affinity chromatography with the biotinylated JMV2959 and homogeneous fractions of GHSR-containing discs finally obtained through a size-exclusion chromatography step on a S200 increase column (10/300 GL) using a 25 mM HEPES, 150 mM NaCl, 0.5 mM EDTA, pH 7.4 buffer as the eluent. The full-length cysteine-free β-arrestin 1 mutant and its pre-activated version (truncation at residue 382) with reactive cysteines at positions 68, 167 or 191 were produced as recently described ^30^. For LRET measurement, a truncated receptor where a sortase ligation consensus sequence (LPERGGH) replaced the 346-366 region was produced using the protocol described above. The receptor in nanodiscs was then incubated overnight at 4°C with 2 μM sortaseA and the peptide GGG-GHSRp where the potential phosphorylation sites in the 346-366 region of GHSR (S349, T350, S362, S363 and T366 ^32^) had been replaced by their phosphorylated counterpart, as described in ^48^. Non-ligated receptor was then removed using a HisTrap 1 mL column (Cytiva).

#### Protein labeling

For labeling β-arrestin 1 with MB (ThermoFisher), the latter was added from a 100 mM stock solution in DMSO to purified arrestin at a 1:10 molar ratio and incubated overnight at 4°C in the dark. The reaction was then stopped with 5 mM L-cysteine and unreacted fluorophore removed on a Zeba Spin column (ThermoFisher). Labeling with the Tb-cryptate for the LRET experiments was carried out by incubating the β-arrestin 1 with 2 equivalents of Lumi-4 Tb maleimide (CisBio) overnight at 4°C, on the one hand, and incubating the GHSR containing a single reactive cysteine at position 71^1.60 29^ with Alexa Fluor 488 maleimide (ThermoFisher) at a 1:1.5 receptor-to-dye molar ratio, at 4°C for 12 hours, on the other hand. The protein samples were then desalted on a Zeba Spin column (ThermoFisher). In all cases, the labeling ratios were calculated from the absorption spectra of the labeled proteins using the known extinction coefficients of the receptor and the fluorophores.

#### MB fluorescence measurements

Fluorescence experiments were performed on a Cary Eclipse spectrofluorimeter (Varian). For each scan, the protein concentration was in the 5 µM range, λexc was set at 380 nm, the excitation and emission bandpass set at 5 nm, and emission measured between 410 nm and 520 nm. Fluorescence intensity was corrected for any dilution effect. For the MB experiments with the probe attached to C191 in β-arrestin 1 and for empty nanodiscs, 10% 16:0 Tempo PC (Avanti polar lipids) were inserted into the nanodiscs ^49^.

#### LRET measurements

LRET was measured with a spectrometer with a pulsed Xe lamp as the excitation source (λexc: 337 nm, λem: 515 nm; 5 µs steps; 100 µs delay for Alexa Fluor 488 λexc: 337 nm, λem: 544 nm; 5 µs steps; 100 µs delay for BodipyTMR). The emission profiles are the mean of three measurements. Donor Tb3+ -chelate fluorescence decays were recorded for each probe position in donor-only labeled complexes. Sensitized emission was fitted to a multi exponential decay function, and the goodness of the exponential fit determined from the random residual. Lifetime distributions 588 shorter than 100 μs were discarded as these are largely a function of the instrument response time ^50^. The slow and fast components correspond to the two major time constants of donor fluorescence decay inferred from these exponentials. The distances between donor and acceptor molecules were calculated from the efficiency of energy transfer ^29^. Molecular fractions were calculated from the pre-exponential factors and the excited state lifetime values ^33^

## Acknowledgements

We thank CNRS, Université de Montpellier, Agence Nationale de la Recherche (ANR-20-CE92-0028, ANR-21-CE29-0012, ANR-22-CE44-0042) and the Fondation Pour la Recherche Médicale (FRM, Equipe FRM EQU202103012736) for their financial support. This work was granted access to the HPC resources of IDRIS under the allocations A0100712444 and A0140714133 made by GENCI ». We thank Dr. Paulo Cesar Telles de Souza for helpful discussions on using the MARTINI3 force-field and to have provided us with PI(4,5)P2 parameters. We are also grateful to the program CAPES-COFECUB (project Ph-C 882-17) which promoted the collaboration between the French and Brazilian teams.

