## Supplementary data for "LIPIDS MODULATE THE DYNAMICS OF GPCR:β-ARRESTIN INTERACTION"

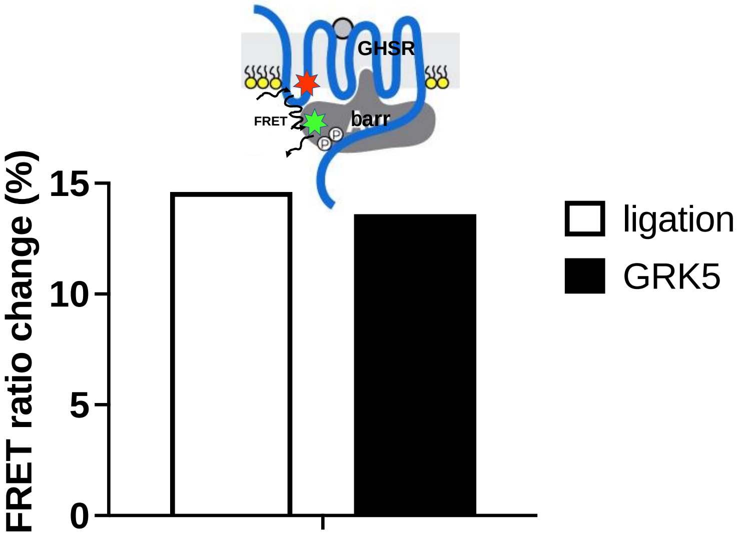


**Supplementary Figure S1: Ligation of a phosphorylated C-tail allows recruitment of the wild-type β-arrestin 1 to the ghrelin receptor.** FRET-monitored wild-type β-arrestin 1 recruitment to MK0677-activated GHSR assembled into MSP1E3D1(-)/POPC/POPG nanodiscs after ligation of a phosphorylated C-tail (ligation) or after *in vitro* phosphorylation with recombinant GRK5 (GRK5). The recruitment assay consisted in measuring the FRET signal between AlexaFluor350 introduced on a unique reactive cysteine of GHSR at position $71^{1.60}$ [1] and AlexaFluor488 introduced on a unique reactive cysteine at position 169 of recombinant β-arrestin 1 [2]. Data are presented as the change in the FRET ratio in the presence of MK0677 compared to that in the absence of ligand. The phosphorylation sites in the GHSR C-terminal region used in the ligation strategy were those previously identified in the literature, *i.e.* S349, T350, S362, S363, and T366 [3].

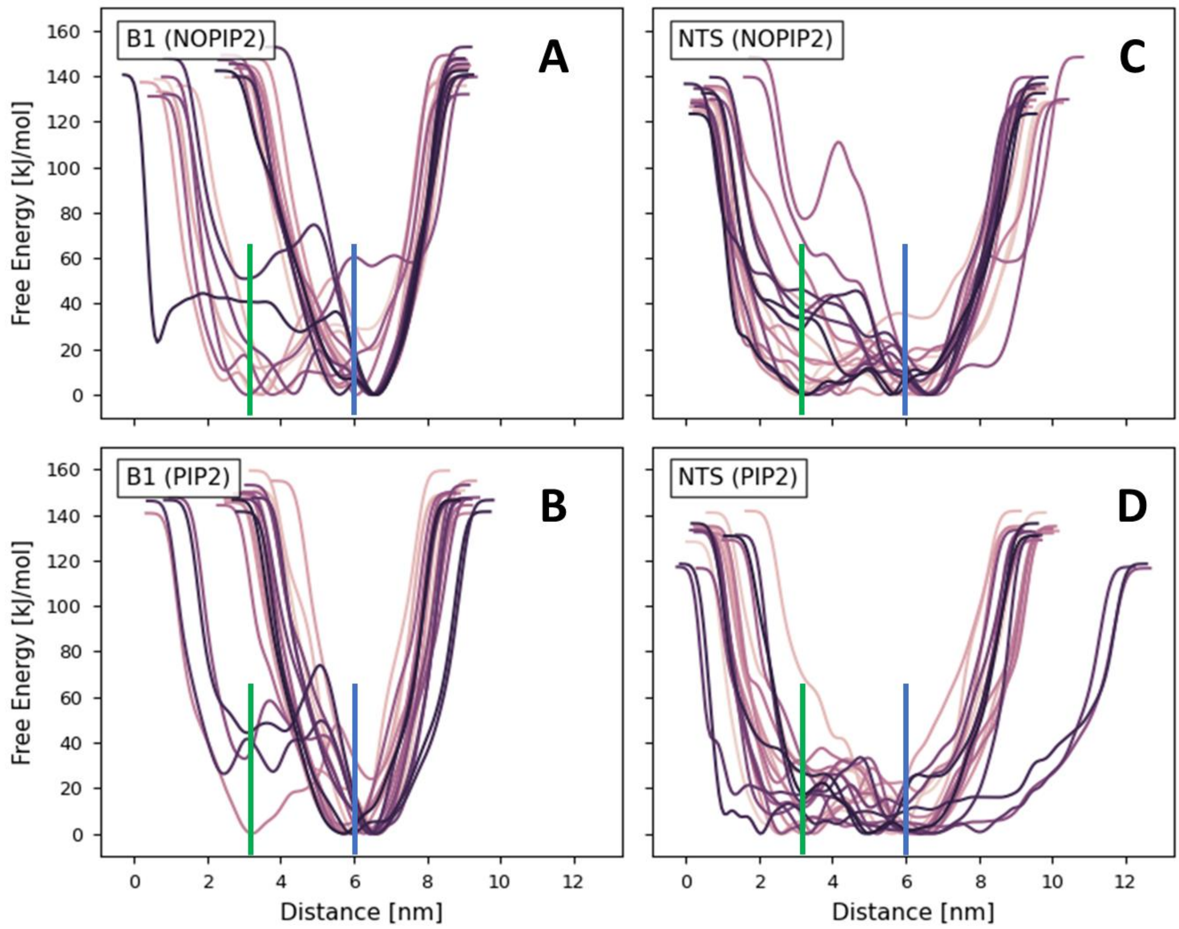


**Supplementary Figure S2**: Free energy surfaces resulting from the meta-dynamics CG simulations of the GHSR: β-arrestin complex. The collective variable here was the distance that was measured experimentally (*e.g.* between residues F71 of the receptor and L191 of arrestin). The starting conformation and membrane composition are indicated on the subplots. The horizontal lines in green and blue depict the distances for the NTS1R and B1AR orientations, respectively.

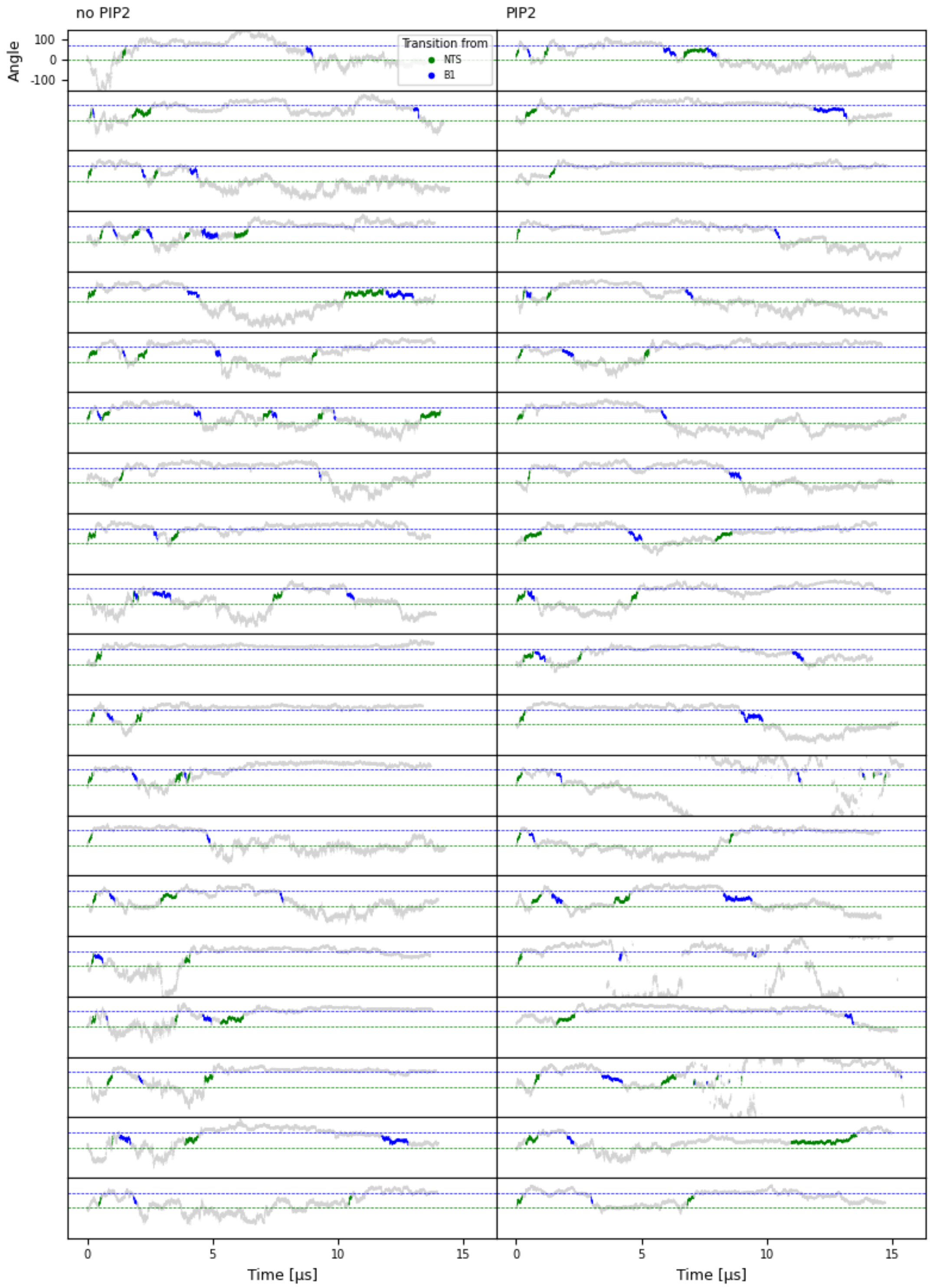


**Supplementary Figure S3 :** Following of the β-arrestin rotation angle in all the CG-MD simulations of the complex starting from the NTS1R orientation. Green and blue regions correspond to transition frames (NTS1R to B1AR in green and B1AR to NTS1R in blue). In three of these simulations, the strong variation of the angle was due to a detachment of β-arrestin from the membrane. These simulations were not conserved in further analyzes.

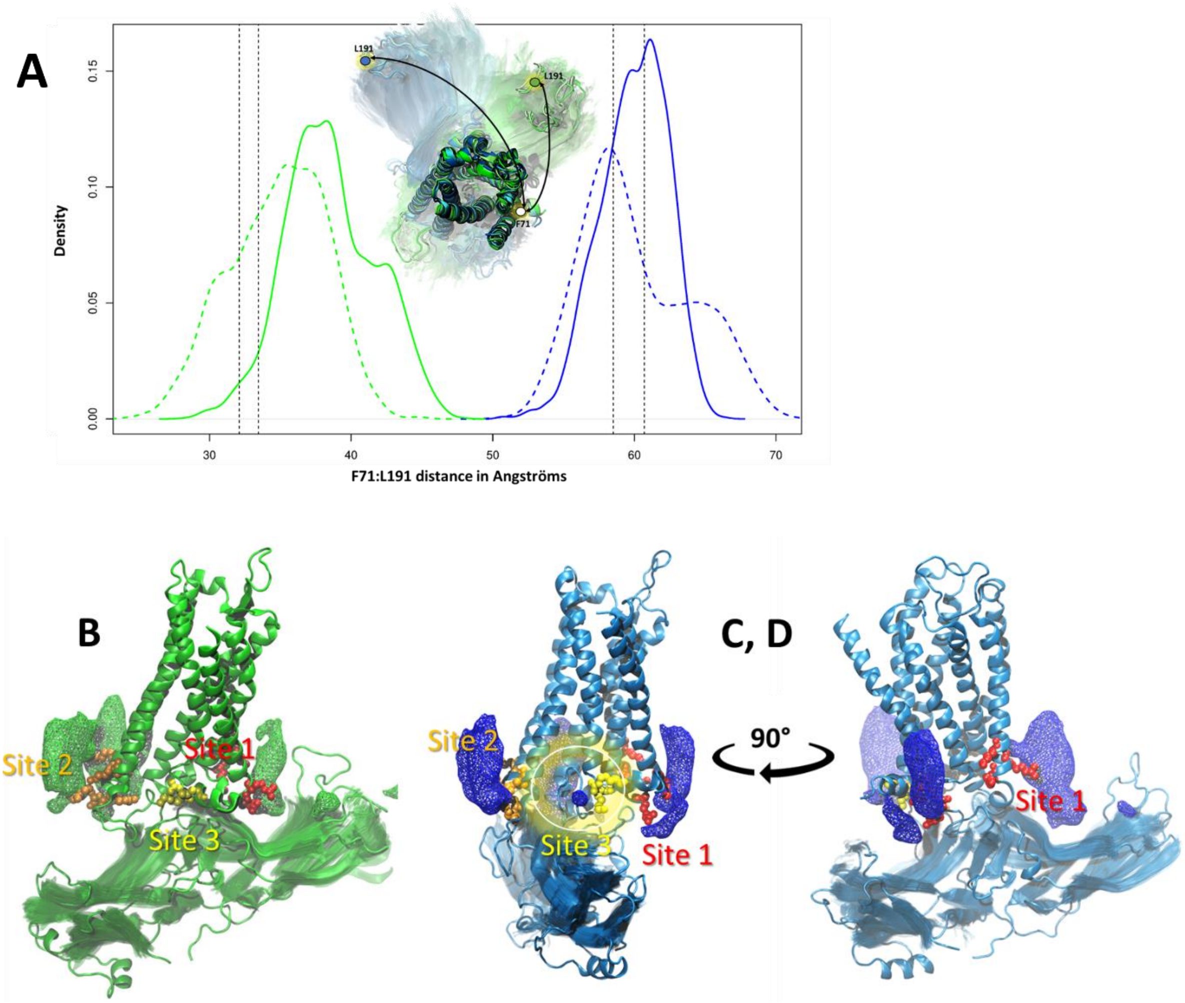


Supplementary Figure S4: Results from all-atoms MD simulations performed on the β-arrestin:GHSR complex and starting with either the NTS1R (in green) or B1AR (in blue) orientations. (A) Distributions of the F71:L191 distance along the obtained trajectories (solid and dashed lines correspond to simulations performed in the presence or absence of PI(4,5)P2, respectively). (B,C,D) Distribution of PI(4,5)P2 around the complex; the three PI(4,5)P2 binding sites we previously identified around the isolated receptor [4] were reported here in red, orange and yellow.
